# Cell division during Xenopus gastrulation influences neuroectoderm patterning

**DOI:** 10.1101/2025.11.11.687238

**Authors:** Ian Velloso, Rodrigo Araujo, Marko Horb, Jose G. Abreu

## Abstract

Oriented cell division is essential for establishing the anterior-posterior body (A-P) axis in diverse species. However, in *Xenopus laevis*, blocking cell division during gastrulation does not impair the development of the neural tube and body elongation. Here, we demonstrate that neither neurulation nor dorsal mesoderm formation is dependent on cell division. On the other hand, neural plate elongation and A-P patterning are impacted in the absence of cell division, resulting in trunk defects and anterior defects that appear during tailbud stages. We also show that cell division is abundant around the ectoderm of the gastrulating embryo. Still, it is more intense in the dorsal ectoderm (presumptive neural plate), where there is a clear preference for A-P oriented cell divisions. Taken together, our results highlight a conserved mechanism of A-P oriented cell division that is present during neural plate elongation and patterning.

## INTRODUCTION

The question of how animals develop their final form has fascinated biologists for centuries. Amphibians’ early embryogenesis is particularly generous in showing the steps by which a round embryo is transformed into a bilaterian elongated body, typical of vertebrates. A process which includes formation and patterning of the notochord, somites, neural plate/tube, and spinal cord (Gilbert and Barresi 2016; Sive, HL; Grainger, RM; Harland 2000). In this context, the concept of neural induction has been extensively explored. The cellular behaviors responsible for forming neural structures, which are aligned with the A-P body axis, vary among vertebrates. In amniotes, for example, convergent extension (CE) is responsible for the elongation of the prospective brain (forebrain, midbrain, and hindbrain) (Hatada and Stern 1994). Whereas the spinal cord is formed through posterior growth, as bipotent neural mesodermal progenitor (NMp) cells proliferate and are allocated either towards dorsal mesoderm or spinal cord (Le Douarin and Halpern 2000; Charrier et al. 1999; Selleck and Stern 1991; Brown and Storey 2000). In amphibians, on the other hand, the whole neural plate is thought to elongate due to convergent extension (CE) of the dorsal marginal zone and the spinal cord is mainly derived from the posterior portion of the neural plate (R. Keller, Shih, and Sater 1992; R. Keller et al. 1992; Sive, HL; Grainger, RM; Harland 2000). Except for the floorplate, which was shown to be assembled through allocation of bipotent cells that lie within the dorsal blastopore lip (López et al. 2005, López et al. 2003). Importantly, however, there is no evidence of proliferating activity of those bipotent cells during gastrulation or neurulation (read Steventon and Arias 2017 for details regarding different mechanisms of spinal cord formation among vertebrates).

The convergent extension phenomenon has been very much explored in *Xenopus laevis* early embryogenesis. It has been established that medio-lateral intercalation, coupled with radial intercalation, of cells composing the dorsal marginal zone (DMZ) of the embryo is a key driving force for CE to take place (R. Keller 1985; R. Keller et al. 1992). At the ventral marginal zone (VMZ), on the other hand, medio-lateral intercalation, uncoupled from radial intercalation, results in thickening, rather than lengthening, of the tissue (R. Keller and Danilchik 1988). It was also proposed that cell division might be important in defining the neural plate territory during gastrulation in *Xenopus laevis* (R. E. Keller 1978), and indeed, cell division occurs with high frequency at the epithelial neuroectoderm cells of the gastrulating *Xenopus laevis* (Chartrain et al. 2011; Hatte et al. 2014; Saka and Smith 2001). Conversely, oriented cell division significantly contributes to the establishment of the anterior-posterior body axis in various organisms, including insects, zebrafish, mice, and chick. (Sander 1976; Concha and Adams 1998; R. Keller 2006; Gong, Mo, and Fraser 2004; G. C. Schoenwolf and Alvarez 1989; Gary C. Schoenwolf and Alvarez 1992; Gary C. Schoenwolf and Yuan 1995; Wei and Mikawa 2000).

Although it has been demonstrated that, in *Xenopus laevis*, late gastrulation and neurulation occur in the absence of cell division (Harris and Hartenstein 1991; Straub, Imhof, and A W Rupp 2021), the embryo does not continue its development if cell division is inhibited from the late blastula stage onward (Harris and Hartenstein 1991). These studies suggest that cell division may be essential for the initial stages of gastrulation. Moreover, it has been demonstrated that blocking cell division during gastrulation and neurulation affects epigenetic control, leading to abnormal histone modification profiles (Straub, Imhof, and A. W. Rupp, 2021). In this study, we have undertaken the challenge of examining the role of cell division during the morphogenesis of the neural plate and dorsal mesoderm, with particular emphasis on the early steps of gastrulation.

## RESULTS

### Blocking cell division during gastrulation, but not afterwards, entails head and trunk defects

To investigate the role of cell division during the morphogenesis of the neural plate and dorsal mesoderm, we employed a combination of two DNA synthesis inhibitors: Hydroxyurea and Aphidicolin (HUA). This established method of inhibiting cell division has already been demonstrated to be effective in *the Xenopus laevis model*. (Harris and Hartenstein 1991; Straub, Imhof, and A W Rupp 2021). Here, we initiated the HUA treatment at stage 9.5 to block cell division from stage 10 to stage 13 (see methods for details). The treatment was confirmed effective, as embryos fixed at stage 13 exhibit many cell divisions when treated with DMSO but show almost none when treated with HUA (Fig. 1D-E). We identified that blocking cell division during this developmental period does not prevent gastrulation and neurulation from occurring (Fig. 1 B and C), consistent with previous studies. Moreover, stage 20 HUA-treated embryos exhibit well-formed notochord, paraxial mesoderm, and neural tube, although they are less robust than DMSO-treated embryos (Fig. 1F, G). Interestingly, we note that embryos with blocked cell division during gastrulation display head defects that are clear at stage 25 (compare Fig. 1H’ with I’). Additionally, they elongate less than DMSO-treated embryos (compare Fig. 1H with I). Later stages reveal severe abnormalities at the anterior end in HUA-treated embryos: underdeveloped heads, a tendency toward cyclopia, and truncated bodies (Fig. 1 J, J’, J’’, K, K’, K’’).

**Figure 1.**
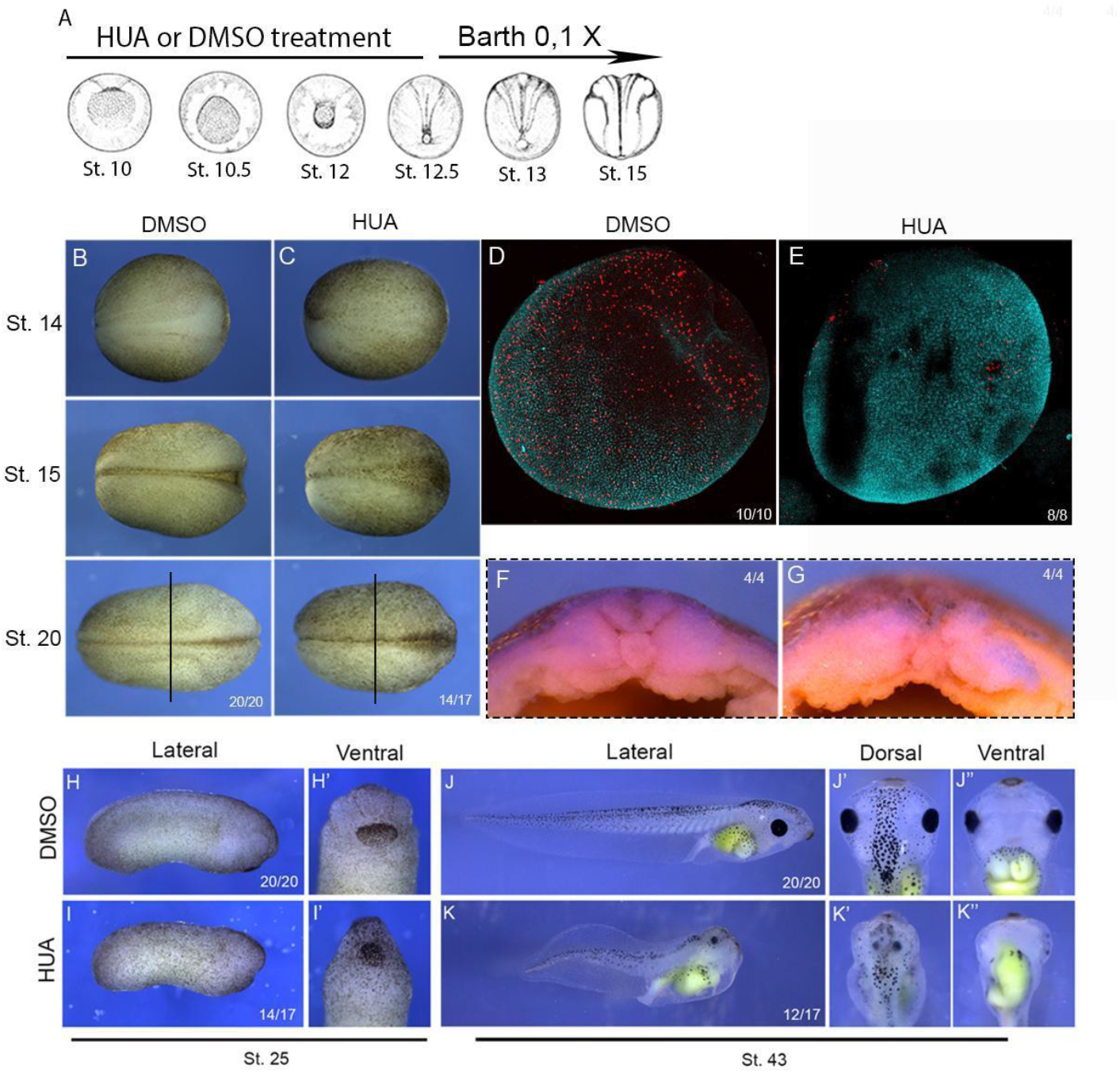
Blocking cell division during gastrulation leads to head and trunk defects. (A) Embryos were treated with a combination of Hydroxyurea and Aphidicolin (HUA) or DMSO from stage 9.5 to stage 12.5 and then transferred to Barth’s 0.1X solution starting at stage 12.5. (B) Dorsal view of a representative embryo treated with DMSO, showing the neural tube closure steps from stage 14 to stage 20. (C) Dorsal view of a representative embryo treated with HUA, showing the neural tube closure steps from stage 14 to stage 20. (D) Dorsal view of an embryo treated with DMSO at stage 13 and immunostained for Histone H2B, showing cells. (E) Dorsal view of an embryo treated with HUA at stage 13 and immunostained for Histone H2B. (F) Transversal section of a DMSO-treated embryo at stage 20, showing notochord, paraxial mesoderm, and neural tube. (E) Transversal section of an HUA-treated embryo at stage 20, showing notochord, paraxial mesoderm, and neural tube. The vertical black line in A and B indicates the site where the transversal section was made. (H) Lateral view of a representative DMSO-treated embryo at stage 25. (H’) Anterior view of a representative DMSO-treated embryo at stage 23. (I) Lateral view of a representative DMSO-treated embryo at stage 25. (I’) Anterior view of a representative HUA-treated embryo at stage 25. (J) Lateral view of a representative DMSO-treated embryo at stage 43. (J’) Anterior dorsal view of a representative DMSO-treated embryo at stage 43. (J’’) Anterior ventral view of a representative DMSO-treated embryo at stage 43. (K) Lateral view of a representative HUA-treated embryo at stage 43. (K’) Anterior dorsal view of a representative HUA-treated embryo at stage 43. (K’’) Anterior ventral view of a representative HUA-treated embryo at stage 43 (stages illustrated according to Nieuwkoop and Faber, 1994).

To determine whether the treatment is stage-specific, we questioned if blocking cell division during neurulation and tailbud stages would also lead to malformations. Remarkably, blocking cell division between stages 13 and 25 did not affect the development of the head or trunk (fig. 2). This confirms that cell division is more crucial for patterning the neural plate before neurulation occurs than during stages when the neural tube is closing, or later during posterior axis elongation (Fig. 2). Having established this, we chose stage 13 (the beginning of neurulation) of development as the ideal moment for a thorough investigation upon the patterning of the dorsal portion of the embryo.

**Figure 2.**
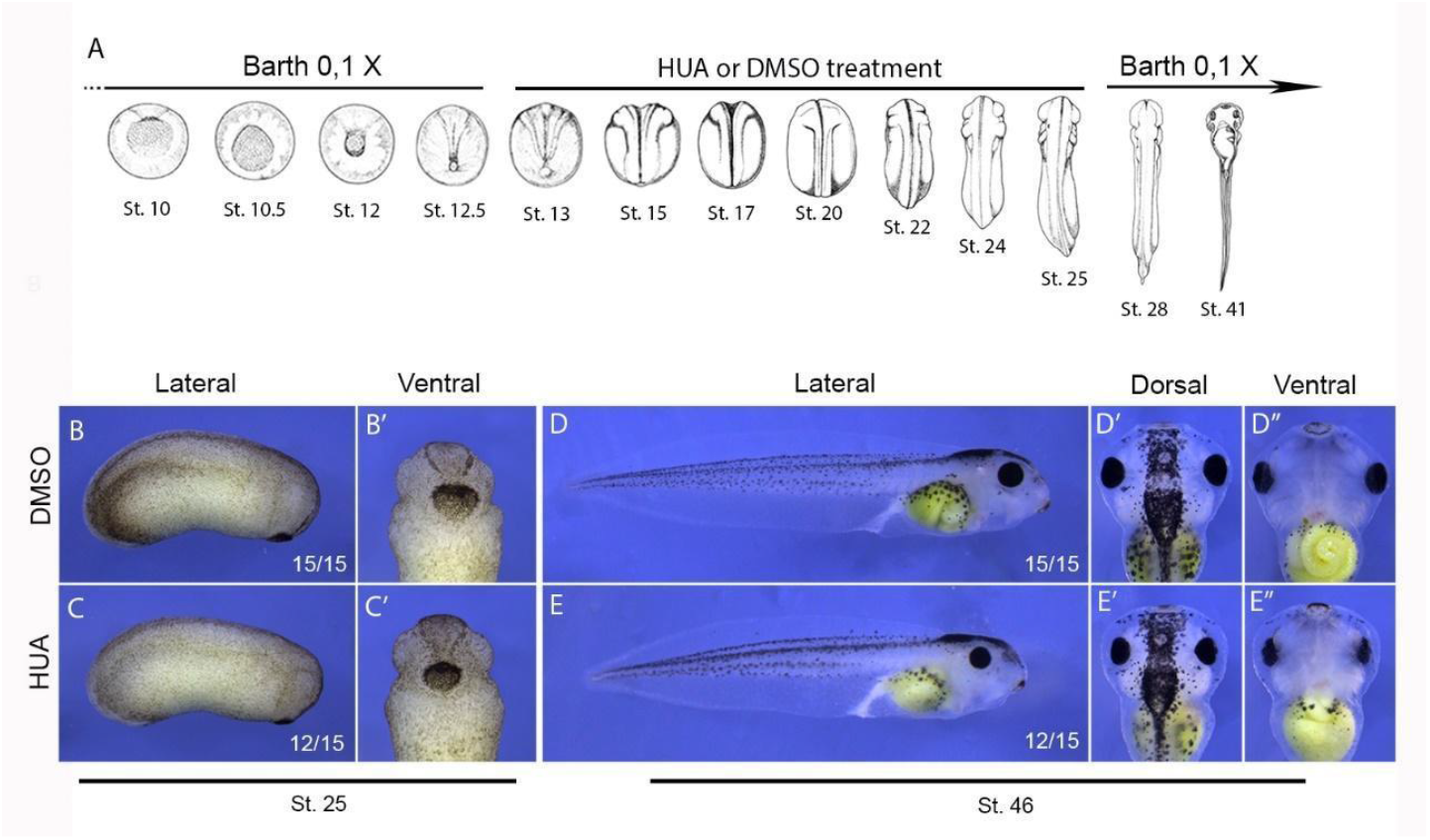
Blocking cell division during neurulation and tailbud stages does not cause head or trunk defects. (A) Embryos were kept in Barth’s 0.1X solution until stage 12.5, then transferred to HUA or DMSO solution, where they remained from stage 12.5 to stage 25, before being returned to Barth’s 0.1X solution. (B) Lateral view of a representative DMSO-treated embryo at stage 25. (B’) Anterior view of a representative DMSO-treated embryo at stage 23. (C) Lateral view of a representative DMSO-treated embryo at stage 25. (C’) Anterior view of a representative HUA-treated embryo at stage 25. (D) Lateral view of a representative DMSO-treated embryo at stage 46. (D’) Anterior dorsal view of a representative DMSO-treated embryo at stage 46. (D’’) Anterior ventral view of a representative DMSO-treated embryo at stage 46. (E) Lateral view of a representative HUA-treated embryo at stage 46. (E’) Anterior dorsal view of a representative HUA-treated embryo at stage 46. (E’’) Anterior ventral view of a representative HUA-treated embryo at stage 46 (stages illustrated according to Nieuwkoop and Faber, 1994).

### Blocking cell division during gastrulation impacts neural patterning in the anterior region

Initially, we conducted a comparative analysis of the neural plate morphological patterns among various embryos treated with either dimethyl sulfoxide (DMSO) or HUA. Although each neural plate exhibits unique characteristics, the DMSO-treated embryos consistently demonstrate a coloration and morphological pattern indicative of healthy embryos: the neural plate region appears lighter than the surrounding ectoderm, and distinct neural folds extend from the anterior end to the blastopore (posterior pole) (Fig. 3a). In contrast, embryos subjected to HUA treatment typically exhibit a significantly reduced area of light ectoderm and markedly discrete neural folds (compare Fig. 3 a and b).

**Figure 3.**
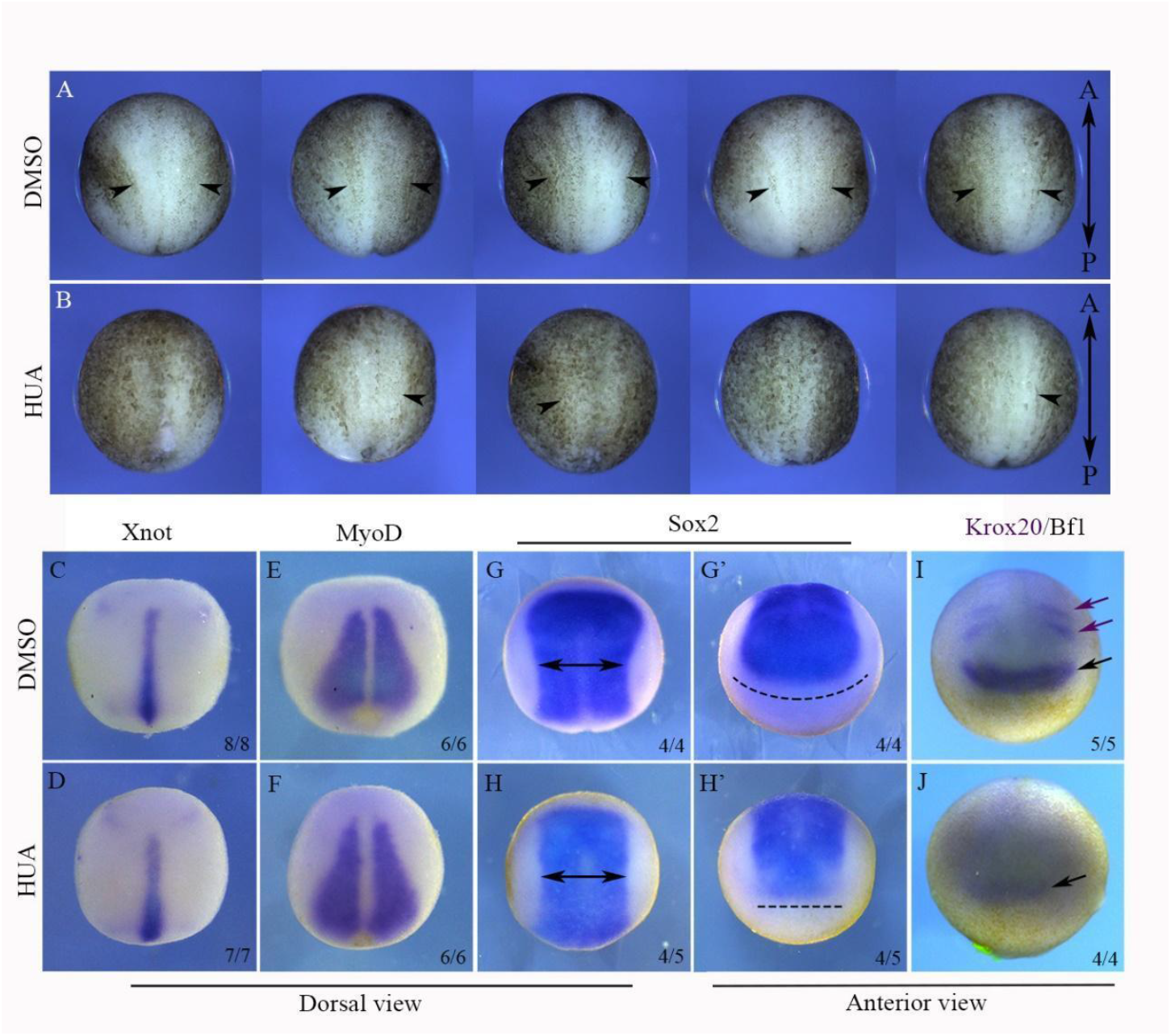
Blocking cell division during gastrulation affects neural plate shaping and patterning. (A) Dorsal view of five embryos at stage 13 treated with DMSO during gastrulation. (B) Dorsal view of five embryos at stage 13 treated with HUA during gastrulation. Arrowheads indicate the lateral hinges of the neural plate. Anterior is always at the top and posterior at the bottom. In situ hybridizations were conducted with embryos at stage 13 treated with DMSO (C, E, G, G’, I) or HUA (D, F, H, H’, J). The embryos were marked for Xnot (C, D), MyoD (E, F), Sox2 (G, G’, H, H’), and Krox20/Bf1 (I, J). Two head arrows in (G) and (H) indicate the Sox2 expression domain width at the most dorsal portion of the embryo, and are the same size. Dashed lines in (G’) and (H’) indicate the width of the most anterior region of the Sox2 domain in each condition. Purple arrows in (I) indicate Krox20 expression domains, while black arrows in (I) and (J) indicate the Bf1 domain.

Having established that HUA-treated embryos display significant differences in neural plate morphological patterning, we next asked if the embryonic territories were altered at stage 13. To address this, we selected marker genes of embryonic territories and performed in situ hybridizations. By comparing *Sox2* domains in both DMSO and HUA embryos, it became evident that blocking cell division impacts the shaping of the neural plate, preventing the full development of its anterior portion (Fig. 3 G, G’, H, and H’). Consequently, anterior markers such as *Bf1* and *engrailed 2* demonstrate a sharp decrease in their expression or are completely lost when cell division is inhibited (Fig. 3 I and J). This suggests that the forebrain, midbrain, and hindbrain domains are not yet patterned at this stage. On the other hand, the axial mesodermal marker *xnot* does not exhibit significant alterations when comparing HUA-treated embryos and DMSO-treated embryos (Fig. 3C and D), indicating that neural plate shaping depends on cell division, whereas notochord shaping does not. Additionally, HUA-treated embryos exhibit a normal pattern of *myoD* expression (Fig. 3E and F), suggesting that paraxial mesoderm morphogenesis is also independent of cell division.

### A-P oriented cell divisions are specifically abundant at the forming neural plate

To understand what is peculiar about the forming neural plate, we decided to track cell movement using transgenic nucleus-GFP embryos - lineage NXR_0139 Xla.Tg (CMV:hist2h2be-GFP;CMV:mRFP). Using low magnification at the light sheet microscope, we were able to detect involution, convergent extension, and epiboly occurring during the formation of the neural plate (supplemental video 1, supplemental figure 1). Remarkably, we could also observe a significant amount of cell division taking place in the ectoderm of the gastrulating embryo (supplemental video 1, supplemental figure 1). Previous cell tracking studies have shown the occurrence of cell division throughout the neural plate during neurulation (Christodoulou and Skourides 2022). Here, we focus on gastrulation and neurulation.

Aiming to determine whether those cell divisions exhibit any specific spatial-temporal pattern, we recorded gastrulation and neurulation with higher magnification and also considered three different angles, in such a way that DNIMZ cells could be tracked alongside lateral non-involuting marginal zone (LNIMZ) and ventral non-involuting marginal zone (VNIMZ) cells (Supplemental Fig. 3 and supplemental videos 2, 3, 4).

All cell divisions were counted as described in the methodology section. Briefly, we identified cells passing through prophase, metaphase, anaphase, and telophase in subsequent frames (usually 3–4 frames, separated by 3 minutes each). By establishing this pattern based on nuclear morphology, we could determine the amount and direction of cell divisions in each recorded area (Supplemental Fig. 2). The recording showed that cell division is present throughout the entire marginal zone of the embryo. Its rate decreases as gastrulation unfolds, taking a significant downward turn as neurulation begins (Fig. 4A). This pattern is somewhat similar all around the marginal zone, but when considering only Anterior-posterior (A-P) oriented cell division, there is an important asymmetry to be highlighted; Only at the DNIMZ, the rate of A-P oriented cell divisions is sustained at the highest rate until stage 12 (Fig. 4D). Later on, it decreases, but not as much as it does in the rest of the marginal zone (Fig. 4D). Indeed, 74% of all divisions happening at the dorsal marginal zone are A-P oriented. In contrast, at the left lateral-ventral region, 66% are A-P oriented. At the right lateral-ventral portion, 55% are A-P oriented (Fig. 4E). Remarkably, this asymmetry gets more drastic when considering only the posterior half of the NIMZ; therein, we have 90% of the DNIMZ cell divisions obeying an A-P oriented direction against 75% and 65% at the left-ventral and right-ventral portions, respectively (Fig. 4F).

**Figure 4.**
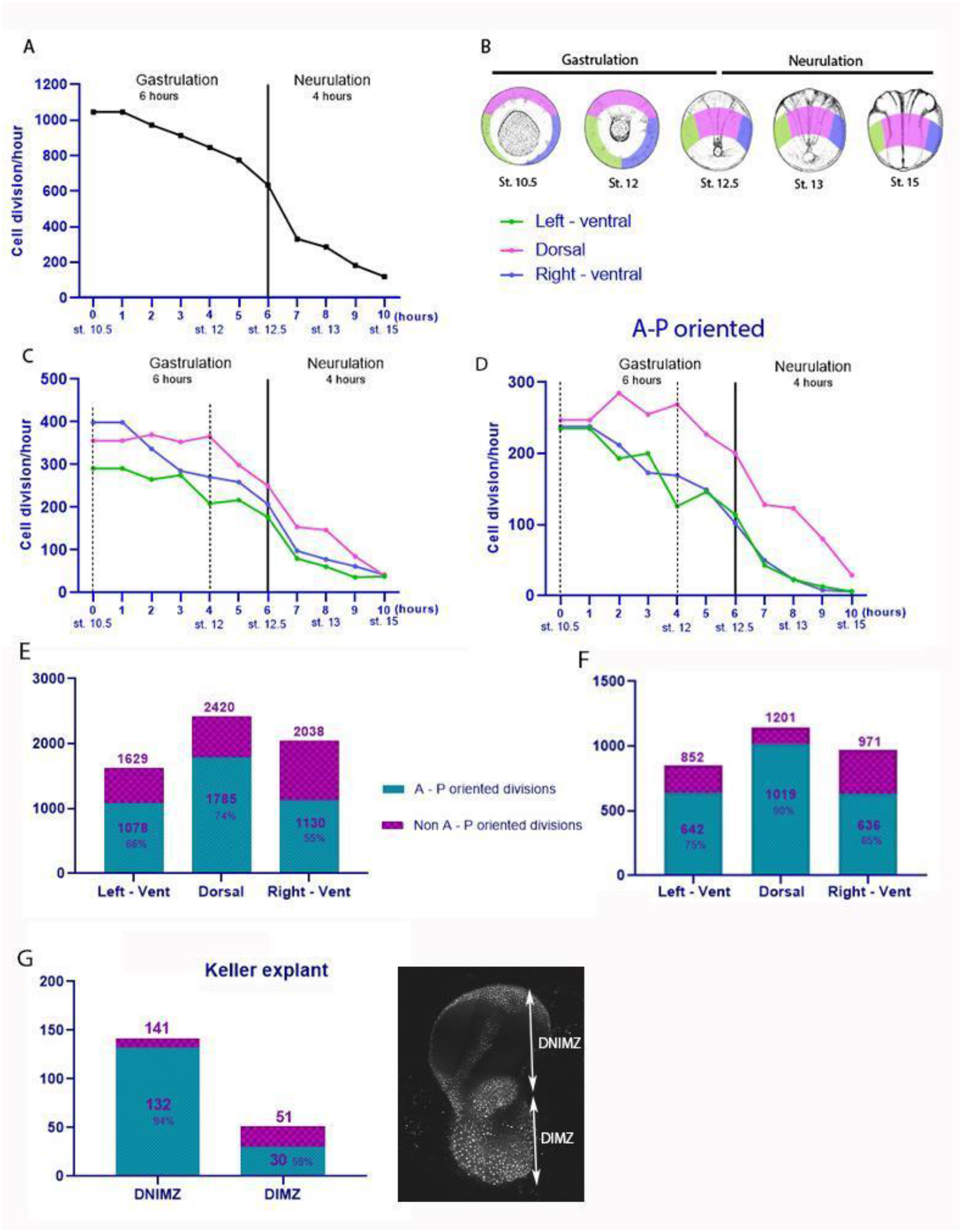
Anterior-posterior (A-P) oriented cell divisions are notably abundant in the developing neural plate. (A) The rate of cell divisions throughout the entirety of the Neural Induction and Migration Zone (NIMZ) during the processes of gastrulation and neurulation. The three distinct areas that were imaged are indicated in panel (B). It is noteworthy that these areas remain consistent as the embryo undergoes transformation throughout gastrulation and neurulation. Pink indicates the dorsal region; green denotes the left-ventral region; and blue represents the right-ventral region. (C) The rate of cell divisions during the stages of gastrulation and neurulation, categorized by areas. (D) The rate of A-P oriented cell divisions during the stages of gastrulation and neurulation, categorized by areas. (E) The total number of cell divisions and the number of A-P oriented cell divisions in each area. (F) The total number of cell divisions and the number of A-P oriented cell divisions counted exclusively in the posterior half of the NIMZ within each area. The graphics from A to F reflect the three time-lapses performed from different angles of the same embryo. (G) Quantification of cell division during the development of the Keller explant at the DIMZ and DNIMZ, which are indicated at a picture of the Keller explant.

We then asked whether this A-P oriented pattern of cell division was specific to the DNIMZ or if the DIMZ also exhibits a similar pattern. To answer this question, we recorded the development of a Keller ‘sandwich’ explant and discovered that DIMZ cells divide significantly less than DNIMZ cells during gastrulation (Fig. 4 G). Additionally, there was no detectable preference for the orientation of these divisions (Supplemental Video 5).

## DISCUSSION

The results presented here demonstrate that cell division is abundant in the superficial layer of gastrulating embryos and tends to diminish as development progresses towards neurulation. Conversely, the inhibition of cell division during the early stages of gastrulation has a significant impact on developmental processes, whereas the obstruction of cell division during the stages of neurulation and tailbud has a negligible effect. We demonstrated that inhibiting cell division during early gastrulation specifically impacts the shaping of the neural plate, which likely explains why anterior neural structures fail to develop properly thereafter. Additionally, we showed that cell division in the neuroectoderm is preferentially oriented along the anterior-posterior axis, and depicting this phenomenon may address an unresolved question related to DMZ lengthening during gastrulation. While it is true that the phenomenon of convergent extension (CE) has been thoroughly examined in *Xenopus laevis*, the reason why early radial intercalation results in anteroposterior (A-P) tissue elongation, rather than an overall spreading in all directions, remains to be elucidated (Wilson and Keller 1991). In discussing this matter, Ray Keller suggests that radial intercalation likely refers to A-P tissue elongation, potentially due to an indiscernible level of medio-lateral intercalation or an unidentified mechanism (Wilson and Keller 1991). Considering the findings presented here, we propose that the oriented cell division of the epithelial dorsal NIMZ is a suitable candidate for causing the tissue to specifically flatten in the A-P direction.

Our findings also indicate a limited occurrence of cell division at the dorsal involuting mesoderm zone (DIMZ) during gastrulation, which aligns with prior observations (Saka and Smith 2001). Indeed, both axial and paraxial mesoderm were demonstrated to develop appropriately in the absence of cell division; this likely explains the well-formed neural tube observed during neurulation. It has recently been shown, through cell tracking experiments, that neural tube closure depends on the Convergent Extension of the posterior neural plate and the apical constriction of the anterior neural plate cells (Christodoulou and Skourides, 2022). Additionally, apart from notochord vertical signaling, mechanical communication from the somitic mesoderm is crucial for neural tube closure to occur (Christodoulou and Skourides 2023).

Indeed, we can conclude from our results that vertical signaling from the axial and paraxial mesoderm partially bypasses early failure in the A-P patterning of the neural plate. Importantly, however, the present study illuminates distinct phases of neural induction and their associated competencies. The initial phase involves the elongation of the neural plate along with anterior-posterior (A-P) patterning, which relies on cellular division. The subsequent phase is the neural tube closure, which is independent of cellular division. Notably, early abnormalities in neural plate patterning do not impair the subsequent neurulation phase; however, they become morphologically apparent at later stages, specifically during the tailbud and larval periods.

It is noteworthy that a significant evolutionary aspect is associated with the distinction between these two phases. While the formation of the neural tube is a mechanism specific to vertebrates, anterior-posterior (A-P) patterning and elongation of the neuroectoderm are embryologically linked to the deeply conserved phenomenon of A-P body axis formation. Indeed, it has been demonstrated that oriented cell division is fundamental for A-P body axis formation across a diverse array of animal species (Sander 1976; Concha and Adams 1998; R. Keller 2006; Gong, Mo, and Fraser 2004; G. C. Schoenwolf and Alvarez 1989; Gary C. Schoenwolf and Alvarez 1992; Gary C. Schoenwolf and Yuan 1995; Wei and Mikawa 2000). In this context, we would like to propose that neural plate elongation and patterning depend on A-P-oriented cell division. Further investigation into the mechanisms responsible for neural plate elongation and patterning has the potential to provide significant insights into the fundamental evolutionary aspects of A-P body axis formation.

## SUPLEMENTAL FIGURES, LEGENDS AND MOVIE LEGENDS

**Figure S1.**
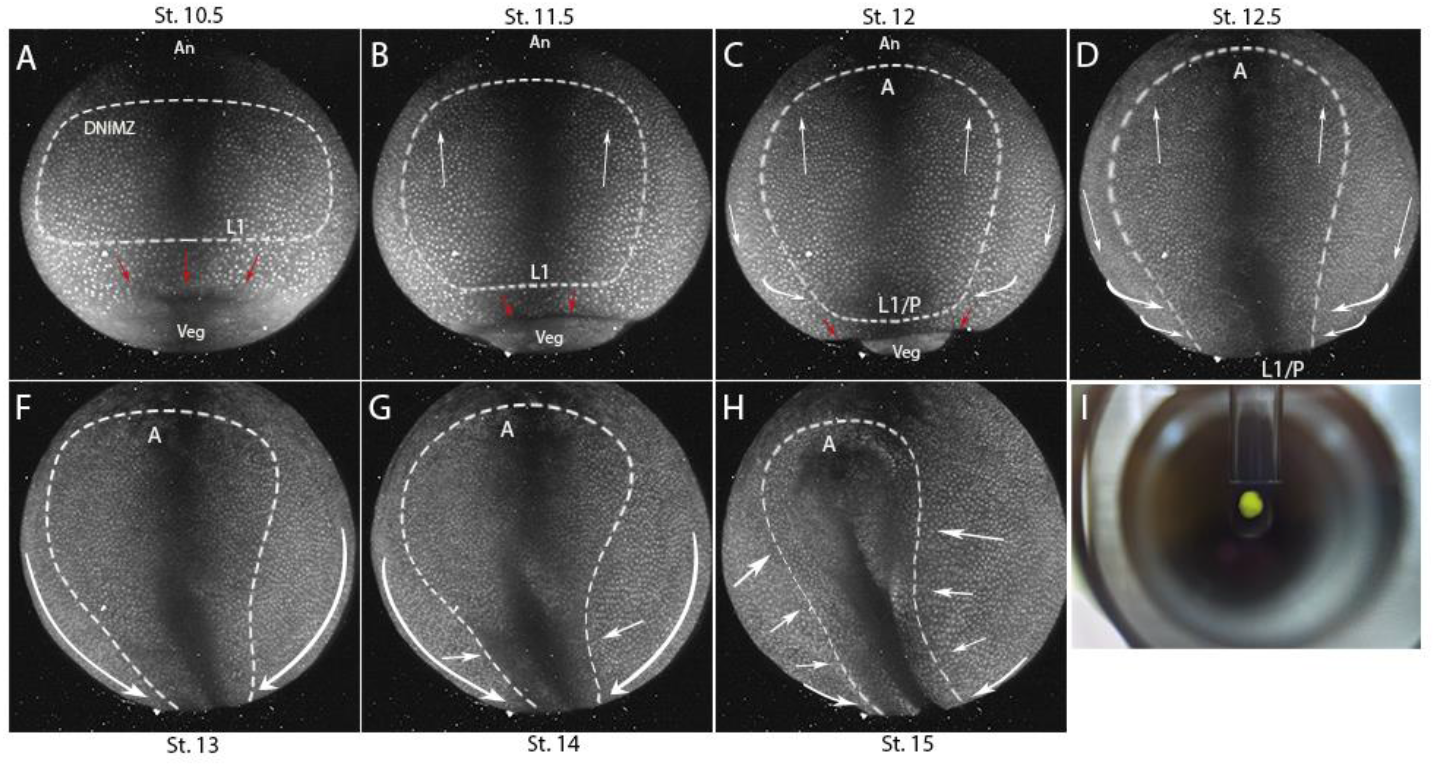
Development of the dorsal NIMZ during gastrulation and neurulation. Time-lapse of a transgenic Xenopus *laevis* embryo for fluorescent nucleus lineage NXR_0139 Xla.Tg(CMV:hist2h2be-GFP;CMV:mRFP, from stage 10.5 (A) of development to stage 15 (H). The approximate space of the DNIMZ is demarcated by a white hatched line. Red arrow indicates movement of cells that make up the DIMZ. White arrows indicate cell movement. Anterior pole (PA). Posterior pole (PP). Approximate border between NIMZ and IMZ (L1). Non-involuting dorsal marginal zone (DNIMZ).(I) *Xenopus laevis* embryo at the gastrula stage mounted in 0.5% agarose inside a glass tube that allows it to fit into the light sheet microscope. This assemblage can be rotated according to the observer’s needs so that the eyepiece, at the back, captures the required angle. The time-lapse was done in a 10X magnification.

**Figure S2.**
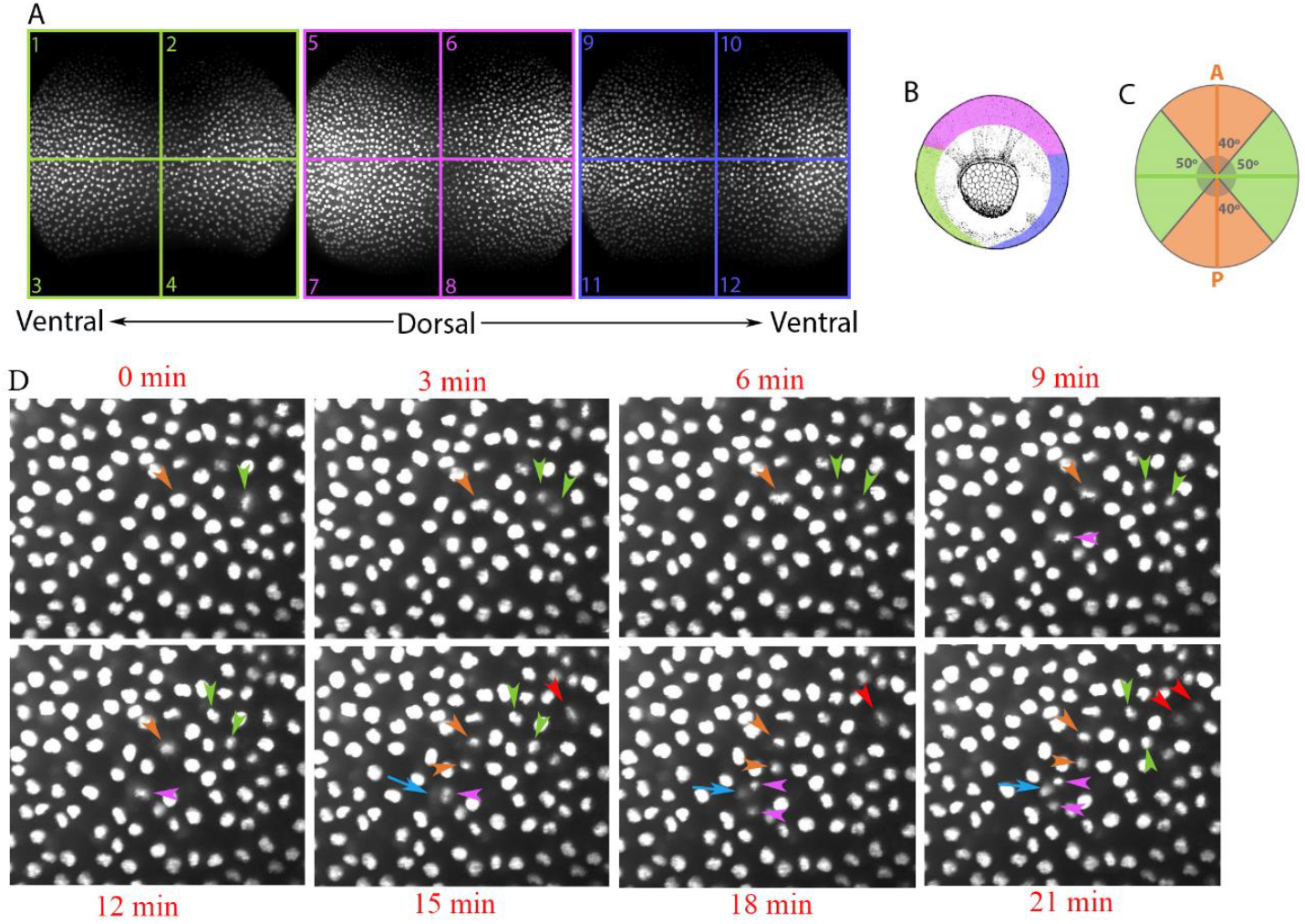
Methodology used to count cell divisions. (A) The three recording angles (green, pink and blue) were each divided into four quadrants. These quadrants (1-12) were used to count the number of cell divisions and the orientation of these cell divisions during the gastrulation and neurulation process. (B) The three recording angles illustrated in the embryo at stage 11.5. (C) Cell divisions considered oriented along the A-P axis are in the orange spectrum (maximum of 40 degrees away from the A-P axis). (D) A 21 minutes period of the time lapse (8 frames out of 200) within the quadrant 7 showing 4 cell divisions taking place. The cell that is about to divide (single arrowheads) shows chromosome alignment before separation (coupled arrowheads). Metaphase is always observable before separation of the chromosomes (note nucleus morphology pointed by green arrowhead at 0 minutes; Orange arrowhead at 6 and 9 minutes; Pink arrowhead at 9 and 12 minutes; and Red arrowhead at 21 minutes). Cell division cannot be mistaken by radial intercalation, which is pointed by the blue arrow. In this case a cell from a deeper layer emerges to the more superficial layer without showing any sign of cell division (note how its signal is almost imperceptible at 15 minutes, then it becomes stronger at 18 minutes and even more at 21 minutes). The case of cell intercalation taking place close to a cell division was specifically chosen to show how those two kinds of events are easy to be distinguished. The images were made in a 20X magnification.

**Figure S3.**
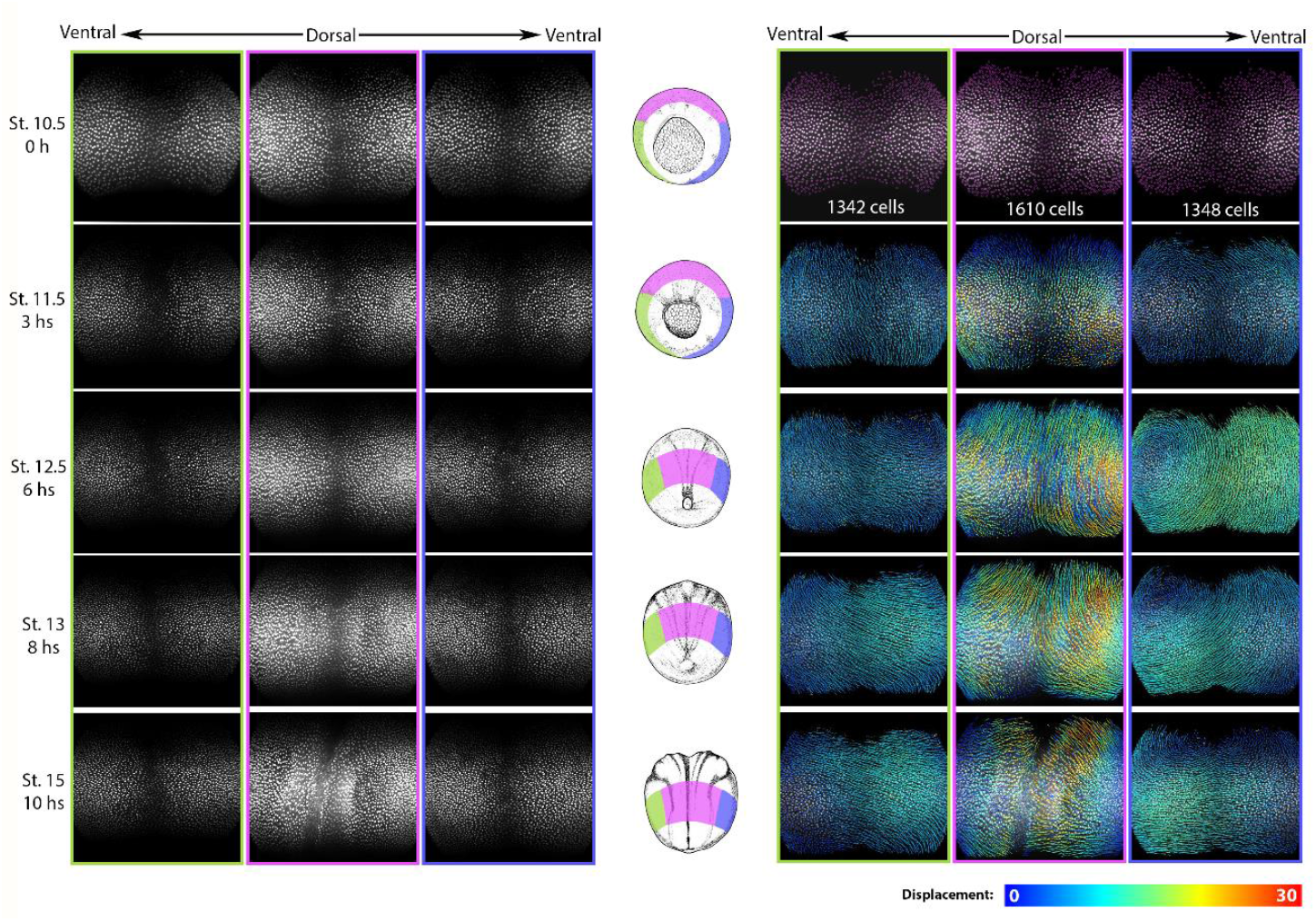
Development of the dorsal, lateral and ventral marginal zone. Time-lapse of a transgenic Xenopus *laevis* embryo for fluorescent nucleus lineage NXR_0139 Xla.Tg(CMV:hist2h2be-GFP;CMV:mRFP, from stage 10.5 to stage 15 of development. One image was made from each angle indicated by the color pattern. Left lateral angle – ventral (green), dorsal angle (pink), right lateral angle – ventral (dark blue). The region imaged for each stage shown is indicated by the color pattern in the middle column (illustrations adapted from *xenbase*.*org*). An analysis performed by track-mate in ImageJ shows us the number of cells present in each of the angles chosen at the beginning of the experiment, and also the displacement route followed by each of these cells. The color pattern indicates the relative displacement of the cells. The red routes present a displacement 30 times greater than the dark blue routes. The images were made in a 20X magnification.

Supplemental video 1. Whole embryo H2b–GFP. From stage 10.5 to stage 15. Dorsal view. Animal pole up, Vegetal pole down. 10X magnification.

Supplemental video 2. Whole embryo H2b–GFP time-lapse. 10 hours of development, 1 frame every three minutes. Focus on dorsal NIMZ. From stage 10.5 to stage 15. Animal pole up, Vegetal pole down. 1 image every 3 minutes. 20 X magnification.

Supplemental video 3. Whole embryo H2b–GFP time-lapse. 10 hours of development, 1 frame every three minutes. Focus on left – ventral NIMZ. From stage 10.5 to stage 15. Animal pole up, Vegetal pole down. 20 X magnification.

Supplemental video 4. Whole embryo H2b–GFP time-lapse. 10 hours of development, 1 frame every three minutes. Focus on right – ventral NIMZ. From stage 10.5 to stage 15. Animal pole up, Vegetal pole down. 20 X magnification.

Supplemental video 5. Keller explant H2b–GFP time-lapse. 10 hours of development, 1 frame every five minutes. From stage 10.5 to stage 15. NIMZ up, IMZ down. 20 X magnification.

## MATERIALS AND METHODS

### Obtaining embryos

Adult frogs (Nasco Inc., WI, USA, or NXR, MBL, MA, USA) were stimulated with human chorionic gonadotropin (HCG-Sigma). *Xenopus laevis* embryos were produced by in vitro fertilization, dejellied with 3% cysteine solution (diluted in Barth 0.1X; pH 7.8), and cultured in Barth 0.1X (Barth 10x: 880 mM NaCl; 10 mM KCl; 10 mM MgSO4; 50 mM HEPES (pH 7.8); 25 mM NaHCO3; Kanamycin 0,1 g). The embryos were staged according to Nieuwkoop and Faber (1994). At NXR, MBL, we used wild-type frogs (NXR_0031) and the transgenic lineage NXR_0139 Xla.Tg(CMV:hist2h2be-GFP;CMV:mRFP).

### Blocking cell division

Cell division was blocked by treatment with the drugs Hydroxyurea and Aphidicolin (HUA). Hydroxyurea inhibits DNA synthesis by blocking ribonucleotide diphosphate reductase, an enzyme that catalyzes the conversion of ribonucleotides into deoxyribonucleotides. This is an essential step in DNA biosynthesis. The drug Aphidicolin blocks eukaryotic DNA polymerase alpha. Aphidicolin stock solution was diluted in DMSO at a concentration of 10 mg/ml and frozen at −20 degrees Celsius in several aliquots. On the day of treatment, the aliquot was thawed and diluted in 0.1X Barth solution at a concentration of 150 μM Aphidicolin and 20 mM hydroxyurea. The control solution was 2% DMSO in 0.1X Brth solution. The embryos used in the experiment were immersed in the HUA or DMSO solution at two different windows of development: 1. From stage 9.5 to stage 12.5 (Fig. 1A); or 2. From stage 13 to stage 25 (Fig. 2A). The concentrations used have already been proven effective in two separate publications, and block cell division after 2-3 hours after the beginning of the treatment (HARRIS, HARTENSTEIN, 1991, STRAUB, IMHOF, et al., 2021). Embryos were kept at 16 °C, so they took approximately 2 hours to progress from stage 9.5 to stage 10.

### In situ hybridization

Whole embryos were fixed in MEMFA 1X (10% de MEM 10X (Mops 1M em pH 7.4, EGTA 20mM, MgSO4 10mM) + 10% formaldehyde 37% + 80% Distilled water) for 2 h at room temperature or at 4°C overnight, and then dehydrated in an ethanol series. Whole-mount in situ hybridization was performed according to Abreu et al. (2002) with modifications suggested by Reversade and De Robertis (2005). After in situ hybridization, the embryos were treated with a bleaching solution (2.5% 20× SSC, 5% formamide, and 4% H2O2 in H2O) and photographed using a digital camera (Leica DFC290 HD) coupled to a microscope (MZ125, Leica).

### Fluorescence Immunocytochemistry

Whole embryos were fixed in MEMFA 1X (10% de MEM 10X (Mops 1M em pH 7.4, EGTA 20 mM, MgSO4 10 mM) + 10% formaldehyde 37% + 80% Distilled water) for 2 h at room temperature or at 4°C overnight and then dehydrated in ethanol series. Whole-mount fluorescence immunocytochemistry was performed according to Lee et al. (2008) using the primary antibody Phospho-Histone H3 (Ser10) (6G3) Mouse mAb (Cell Signaling 9706) and the secondary antibody Alexa Fluor 546 goat anti-mouse IgG (REF A11003). After immunocytochemistry, embryos were imaged using a stereomicroscope (MZ10F, Leica).

### Nucleus-GFP time-lapse imaging

To perform the Time-lapse at a cellular level, the Zeiss light-sheet 7 microscope, installed in the Central Microscopy Facility at Marine Biological Laboratory, Woods Hole, United States, was used. The microscope was programmed to take a complete image of the embryo or explant every three minutes for a total period of 10 hours. The nucleus-GFP embryos were obtained from the transgenic lineage NXR_0139 Xla.Tg (CMV:hist2h2be-GFP; CMV:mRFP) were excited with a laser emitting a wavelength of 488 nm. The embryos were mounted inside a plastic tube so that it could be rotated 120 degrees, and thus the entire extension of the embryo’s DMZ could be photographed every three minutes. The acquisitions were performed with 20x objectives and with a 3 μm increment between each photographed plane. Importantly, the distinct focus planes are capable of capturing different heights of the ectoderm, but deep layers are not captured.

To quantify all cell divisions, we divided each of the time-lapses into four quadrants (Supplemental Fig. 2), thus generating twelve distinct quadrants. Each of the quadrants was analyzed frame by frame (10 hours of recording captured by 200 frames, with a 3-minute increment). It was possible to identify a morphological pattern typical of cells during their division cycle: chromosomes aligning at the cell’s equator (i.e., metaphase), followed by the gradual separation of chromosomes (i.e., anaphase – telophase). Cell divisions occurring within a 40-degree angle from the A-P axis were classified as A-P oriented. Cell intercalation events were also observed, but could be easily distinguished from cell division because they display a different pattern: a gradual increase in nuclear signal as the cell emerges to the surface of the embryo, with no change in nuclear morphology (Supplemental fig. 2 and supplemental videos 1, 2, and 3).

### Keller explant

All microsurgical procedures were performed using a magnifying stereo microscope (Leica S8APO) and tools fabricated by the experimenter, including a hair-looping instrument and a hair knife. The Keller explant technique, widely used in embryological studies, aims to examine cell movements involved in the gastrulation of *Xenopus laevis* (DONIACH, PHILLIPS, et al., 1992, KELLER, Ray, TIBBETTS, 1989, WILSON, KELLER, 1991). It is the cultivation of the dorsal marginal region of an embryo adjacent to the dorsal marginal region of another embryo. Both explanted at the early stage of gastrulation (st. 10) and connected as a sandwich that preserves the deep cells inside and the epithelial cells outside. The explant should be composed of the involuting marginal zone (IMZ): deep cells destined to become notochord, somites, and an endodermal epithelial layer; and the non-involuting marginal zone (NIMZ): ectodermal cells. The cell group committed to actively migrating to form the prechordal mesendoderm should be carefully removed from the explant.

## Acknowledgements

We acknowledge funding from CNPQ (313023/2020-4 and 313588/2025-2), funding from FAPERJ (E-26/200.936/2022, E-26/010.000080/2018 and SEI-260003/010293/2023.) We acknowledge funding from National Institute of Health (NIH R24OD030008 and P40OD010997). We acknowledge Dr. Carsten Wolf, who supervised every experiment in the Central Microscopy Facility at the Marine Biological Laboratory, and helped us to create the protocol of imaging the fluorescent embryos.

